# Evolution of *lbx* spinal cord expression and function

**DOI:** 10.1101/2020.12.30.424885

**Authors:** José Luis Juárez-Morales, Frida Weierud, Samantha J. England, Celia Demby, Nicole Santos, Ginny Grieb, Sylvie Mazan, Katharine E. Lewis

**Affiliations:** Department of Biology, Syracuse University, 107 College Place, Syracuse, NY 13244, USA; Programa de Cátedras CONACyT, Laboratorio de Patogénesis Microbiana, Centro de InvestigacionesBiológicas del Noroeste, S.C.(CIBNOR), La Paz, Baja California Sur, México; Department of Physiology, Development and Neuroscience, University of Cambridge, Downing Street, Cambridge CB2 3DY, UK; Biologie Intégrative des Organismes Marins, UMR 7232 CNRS, Observatoire Océanologique, Sorbonne Université, Banyuls-sur-Mer, France

**Author notes:** Corresponding author. Department of Biology, Syracuse University, 107 College Place, Syracuse, NY 13244, USA. Ethics Approval: All zebrafish experiments were approved by UK Home Office or Syracuse University IACUC committee.

**Keywords:** dI4, dI5, dI6, small-spotted catshark, shark

## Abstract

Ladybird homeobox (Lbx) transcription factors have crucial functions in muscle and nervous system development in many animals. Amniotes have two *Lbx* genes, but only *Lbx1* is expressed in spinal cord. In contrast, teleosts have three *lbx* genes and we show here that zebrafish *lbx1a*, *lbx1b* and *lbx2* are expressed by distinct spinal cell types, and that *lbx1a* is expressed in dI4, dI5 and dI6 interneurons, as in amniotes. Our data examining *lbx* expression in *Scyliorhinus canicula* and *Xenopus tropicalis* suggest that the spinal interneuron expression of zebrafish *lbx1a* is ancestral, whereas *lbx1b* has acquired a new expression pattern in spinal cord progenitor cells. *lbx2* spinal expression was probably acquired in the ray-finned lineage, as this gene is not expressed in the spinal cords of either amniotes or *S. canicula*. We also show that the spinal function of zebrafish *lbx1a* is conserved with mouse Lbx1. In zebrafish *lbx1a* mutants, there is a reduction in the number of inhibitory spinal interneurons and an increase in the number of excitatory spinal interneurons, similar to mouse *Lbx1* mutants. Interestingly, the number of inhibitory spinal interneurons is also reduced in *lbx1b* mutants, although in this case the number of excitatory interneurons is not increased. *lbx1a;lbx1b* double mutants have a similar spinal interneuron phenotype to *lbx1a* single mutants. Taken together these data suggest that *lbx1b* and *lbx1a* may be required in succession for correct specification of dI4 and dI6 spinal interneurons, although only *lbx1a* is required for suppression of excitatory fates in these cells.

**Research Highlights:** *lbx1* spinal expression and function is conserved in vertebrates. In contrast, zebrafish *lbx1b* and *lbx2* have novel spinal expression patterns that probably evolved in the ray-finned vertebrate lineage (*lbx2*) or teleosts (*lbx1b*).

## Introduction

The spinal cord is a crucial part of the central nervous system of all vertebrates as its neuronal circuitry controls movements and receives sensory inputs from the trunk and limbs. All of the data so far, suggest that spinal cord patterning and neuronal circuitry are highly conserved in extant vertebrates, although in amniotes, some populations of spinal neurons have diversified into specialized sub-classes of highly-related neurons. These data also suggest that the common ancestor of ray-finned and lobe-finned vertebrates had distinct classes of spinal neurons with particular functions, that were specified during embryonic development by different transcription factors (e.g. Alvarez et al., 2005; Goulding & Pfaff, 2005; Griener et al., 2015; Lewis, 2006). For example, analyses by ourselves and others have identified several transcription factors that are expressed at conserved dorsal-ventral positions in both amniote and zebrafish spinal cord, although the expression domains are usually larger in amniotes, corresponding to the larger spinal cords in these vertebrates (e.g. Batista et al., 2008; Batista & Lewis, 2008; Juárez-Morales et al., 2016; Moran-Rivard et al., 2001). Consistent with these conserved expression patterns, the types of neurons found in the spinal cord, and the functions of particular transcription factors in specifying these neuronal subtypes is also highly conserved in different vertebrates (e.g. Goulding & Pfaff, 2005; Juárez-Morales et al., 2016; Lewis, 2006). However, most of these comparative analyses have so far been limited to the ventral spinal cord and it is still unclear whether dorsal spinal neurons are as highly conserved.

The ventral spinal cord primarily contains neurons that are involved in controlling movement and relaying information about trunk and limb position. In contrast, the dorsal spinal cord primarily contains neurons that process and relay sensory information. We already know that there is at least one difference between the neurons in amniote and anamniote dorsal spinal cords, as anamniote embryos have a transient population of large sensory neurons, called Rohon-Beard (RB) cells, that form in the most dorsal part of the spinal cord, whereas amniote embryos do not ((Lewis & Eisen, 2003) although see (Reyes et al., 2004) for reports of possible amniote RB-like cells). However, these RB cells are lost during development and their functions are subsumed by dorsal root ganglia neurons, which are sensory neurons in the peripheral nervous system, that exist in both amniote and anamniotes (Reyes et al., 2004). Both amniote and anamniote embryos have dorsal spinal interneurons, although as mentioned above, the extent to which the specification and/or functions of these interneurons are conserved between different vertebrates is still unclear. This is an interesting and important question from both an evolutionary perspective and also for evaluating the efficacy of different animals as model systems for human spinal cord.

Ladybird homeobox (Lbx) transcription factors have crucial functions in muscle development in many different animals (e.g. Brohmann et al., 2000; Gross et al., 2000; Lou et al., 2012). In addition, they are required for specification of particular cells in the Drosophila nervous system and mammalian spinal cord (Cheng et al., 2005; De Graeve et al., 2004; Gross et al., 2002; Kruger et al., 2002; Müller et al., 2002). All vertebrates examined so far, except teleosts, have two distinct *Lbx* genes, *Lbx1* and *Lbx2* (Wotton et al., 2008; Wotton et al., 2010). In contrast, teleosts have 3 *lbx* genes: *lbx1a*, *lbx1b* and *lbx2* (Wotton et al., 2008; Wotton et al., 2010). In mouse, Lbx1 is expressed in three early-forming (dI4, dI5 & dI6) and two later-forming (DIL_A_ & DIL_B_) classes of post-mitotic dorsal spinal interneurons and it is essential for correct specification of these interneurons (Cheng et al., 2005; Gross et al., 2002; Kruger et al., 2002; Müller et al., 2002). Spinal cord expression of chick Lbx1 is very similar (Schubert et al., 2001). However, *Lbx2* is not expressed in the amniote spinal cord (Chen et al., 2001; Chen et al., 1999; Kanamoto et al., 2006). In contrast, our preliminary data and results from other labs suggest that all three teleost *lbx* genes are expressed in spinal cord (Lukowski et al., 2011; Neyt et al., 2000; Ochi & Westerfield, 2009; Wotton et al., 2008). However, crucially, the specific spinal cord cells types that express each of these genes has not been previously identified. In this paper, we confirm that in contrast to amniotes, not only are all three zebrafish *lbx* genes expressed in spinal cord but their expression domains are distinct from each other. Our data suggest that zebrafish *lbx1a*, like mouse *Lbx1*, is expressed by dI4, dI5 and dI6 spinal interneurons. In contrast, zebrafish *lbx1b* is expressed by spinal cord progenitor cells in the dP4 and potentially also the dP5 domain, whereas zebrafish *lbx2* is expressed in two distinct spinal cord domains, progenitor cells which are probably located in the p1 domain, and late progenitor / early post-mitotic cells in the dI4-dI6 domain.

To address where these differences in *Lbx* spinal cord expression evolved, we examined *lbx* expression in an anamniote tetrapod *Xenopus tropicalis* and a shark, *Scyliorhinus canicula* (also known as small-spotted catshark). Our results show that spinal expression of *lbx1* in *X. tropicalis* and *S. canicula* closely resembles spinal expression of *Lbx1* in mouse and chick and *lbx1a* in zebrafish, suggesting that this expression pattern is ancestral. In contrast, *lbx2* is not expressed in *S. canicula* spinal cord, suggesting that the spinal expression of *lbx2* that exists in zebrafish was probably acquired in the ray-finned vertebrate lineage. As spinal expression of zebrafish *lbx1b* also differs from Lbx1 expression in any of the other vertebrates examined so far, it is likely that this expression pattern was acquired in teleosts, after the duplication of *lbx1* into *lbx1a* and *lbx1b* (Wotton et al., 2008; Wotton et al., 2010). Consistent with similarities between the spinal expression of *lbx1a* in zebrafish and *Lbx1* in mouse, we also demonstrate that zebrafish *lbx1a*, like mouse *Lbx1*, is required for correct specification of a subset of dorsal spinal interneurons. In zebrafish *lbx1a* mutants there is a reduction in the number of inhibitory interneurons and an increase in the number of excitatory interneurons, just like in mouse *Lbx1* mutants (Cheng et al., 2005; Gross et al., 2002; Müller et al., 2002). Interestingly, we also see a reduction of inhibitory spinal interneurons in *lbx1b* mutants, although in this case the number of excitatory interneurons is not increased and *lbx1a;lbx1b* double mutants do not have a more severe spinal cord phenotype than *lbx1a* single mutants. These data suggest that *lbx1b* and *lbx1a* are required, in succession, for specification of inhibitory fates, although only *lbx1a* is required for suppression of excitatory fates, in dI4 and dI6 interneurons. Taken together, our findings identify novel spinal cord expression patterns for zebrafish *lbx1b* and *lbx2*, while also demonstrating evolutionary conservation of *Lbx1/lbx1a* spinal cord expression and function between zebrafish and amniotes, suggesting that the specification of at least some dorsal spinal neurons is conserved between these vertebrates.

## Material and methods

### Zebrafish husbandry and fish lines

All zebrafish experiments were approved by UK Home Office or Syracuse University IACUC committee. Zebrafish (*Danio rerio*) were maintained on 14 hour light/10 hour dark cycle at 28.5°C. Zebrafish embryos were obtained from natural, paired and/or grouped spawnings of wild-type (WT; AB, TL or AB/TL hybrids), *Tg(evx1:EGFP)^SU1^* (Hilinski et al., 2016), *smoothened^b641^* (Varga et al., 2001), *lbx1a^hu3569^*, *lbx1a^sa1496^*, *lbx1b^hu3534^*, (Kettleborough et al., 2013), *mindbomb1^ta52b^* (*mib1)* (Jiang et al., 1996), *Tg(lbx1b:EGFP)^ua1001^* (Lukowski et al., 2011), *Tg(0.9 lbx1a:EGFP)^SU32^* or *Tg(1.6 lbx1a:EGFP)^SU33^* adults. Embryos were reared at 28.5°C and staged by hours post fertilization (h) and/or prim staging as in (Kimmel et al., 1995).

*lbx1a^hu3569^*, *lbx1a^sa1496^* and *lbx1b^hu3534^* mutant alleles were obtained from the Wellcome Trust Sanger Center, (https://www.sanger.ac.uk/resources/zebrafish/zmp/#t_about) (Kettleborough et al., 2013). Each mutation is a single base pair change (C to T) that results in an immediate premature stop codon. In the case of *lbx1a^hu3569^* and *lbx1a^sa1496^*, the stop codon is located 148 bp and 34 bp after the beginning of the homeobox respectively. In *lbx1b^hu3534^*, the stop codon is located 145 bp before the homeobox. Therefore, if truncated mutant proteins are made, Lbx1b^hu3534^ will lack all, Lbx1a^sa1496^ will lack almost all and Lbx1a^hu3569^ will lack part of the C- terminal part of the homeobox domain.

### Creation of *Tg(0.9 lbx1a:EGFP)^SU32^* and *Tg(1.6 lbx1a:EGFP)^SU33^* transgenic lines

Potential *lbx1a* enhancer regions were identified by multispecies comparisons using Shuffle-LAGAN (Brudno et al., 2003) and visualized using VISTA (Mayor et al., 2000). Zebrafish (*Danio rerio*) *lbx1a* (ENSDARG00000018321, Zv9), *lbx1b* (ENSDARG00000018611, Zv9) and orthologous sequences from human (ENSG00000138136, NCBI36 Ensembl release 54) and mouse (ENSMUSG00000025216, NCBIM37 Ensembl release67) were obtained from Ensembl (http://www.ensembl.org). The *Scyliorhinus canicula lbx1* (NC_052161.1) sequence was obtained from https://www.ncbi.nlm.nih.gov/Taxonomy/Browser/wwwtax.cgi?id=7830. *Danio rerio lbx1a* sequence was used as baseline and annotated using exon/intron information from Ensembl. The alignment was performed using 100 bp window and cutoff of 70 % identity. A comparison of approximately 22Kb of *Danio rerio* genomic sequence extending 10 Kb either side of *lbx1a* identified two Conserved Non-coding Elements (CNEs) located 3’ to *lbx1a*. The first is 1037 bp downstream of the stop codon and is 204 bp long whereas the second is 3021 bp downstream of the stop codon and extends over 1060 bp (Fig. 1). Using genomic DNA, we PCR-amplified an amplicon of 900bp around the CNE closest to the 3’ end of *lbx1a* using the following primers, Forward: GTATGCCTGTAAGTGCC, Reverse: CCATCCATAGTGTGACT. We also amplified the 1686 bp CNE using primers Forward: CTTCGTCGCAACTATGA and Reverse: TATTAGCCCAGTAATCA. PCR conditions were: 98°C for 30 s followed by 30 cycles of 98°C 10 s, 62°C 30 s, 72°C 30 s and a final extension step of 72°C for 10 min.

**Figure 1.**
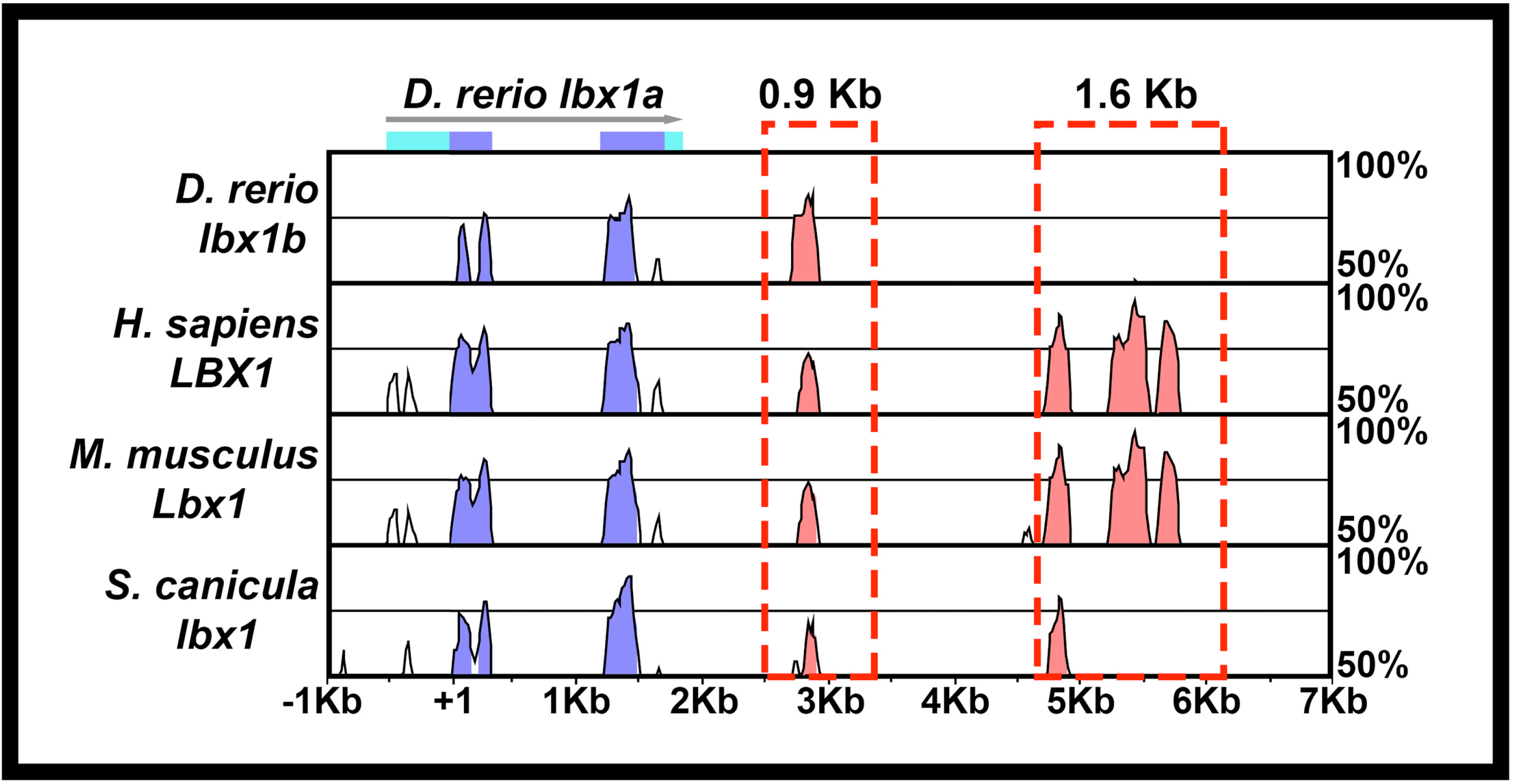
Construction of *Tg (0.9 lbx1a:EGFP)^SU32^* and *Tg(1.6 lbx1a:EGFP)^SU33^* transgenic lines. Schematic showing Shuffle-MLAGAN analysis of *Danio rerio lbx1a* genomic region with zebrafish sequence as baseline compared to *Danio rerio lbx1b* genomic sequence and orthologous regions in *mouse and humanHomo sapiens, Mus musculus* and *Scyliorhinus canicula* genomes. Conserved coding sequences are indicated in blue, arrow indicates 5’→3’ orientation, light blue boxes indicate untranslated regions of *D. rerio lbx1a*. Conserved non-coding elements (CNEs) in 3’ region are indicated in pink. The 0.9 Kb and 1.6 Kb regions amplified to create the *Tg(0.9 lbx1a:EGFP)^SU32^* and *Tg(1.6 lbx1a:EGFP)^SU33^* transgenic lines areis indicated with a red dotted boxes.

Separate reporter constructs were generated for each of the two *lbx1a* CNEs. First, the 900 bp and 1.6 Kb amplicons were cloned into the pDONR™ P4-P1R vector from Invitrogen using Gateway technology (Sasaki et al., 2004; Suzuki et al., 2005). One construct was assembled using the 900 bp *lbx1a* 5′ pDONR with the *cfos minimal promoter:Gal4VP16:UAS:EGFP* middle entry construct (Juárez-Morales et al., 2016; Koster & Fraser, 2001) and the pCSDest2 vector to generate *Tg(Tol2:900bp3’zfish_lbx1a:cfos minimal promoter:Gal4VP16:UAS:EGFP:pA:Tol2)*. For the second construct, the *cfos minimal promoter:Gal4VP16;UAS:EGFP* middle entry vector was modified by removing the *Gal4VP16;UAS* amplification cassette. The final construct was generated using the 1.6 Kb *lbx1a* 5′ pDONR, the *cfos minimal promoter:EGFP* middle entry vector and the pCSDest2 vector (Villefranc et al., 2007). This resulted in a vector containing *Tol2:1.6 Kb 3’ zfish lbx1a:cfos minimal promoter:EGFP:Tol2*.

Plasmid DNA and *transposase* mRNA for microinjection was prepared as in (Juarez-Morales et al., 2017; Kwan et al., 2007). Approximately 10 nl of a combination of plasmid DNA [60-80 ng/μL] and *transposase* mRNA [30 ng/μL] was injected into both blastomeres of 1-2-cell stage zebrafish embryos. Embryos were raised to adulthood and out-crossed to identify founders to generate stable *Tg(0.9 lbx1a:cfos:Gal4VP16;UAS:EGFP)^SU32^* and *Tg(1.6 lbx1a:cfos:EGFP)^SU33^* lines, which we refer to as *Tg(0.9 lbx1a:EGFP)^SU32^* and *Tg(1.6 lbx1a:EGFP)^SU33^*.

### Genotyping

Genotyping was performed on live adults and fixed embryos using DNA from fin clips and dissected heads respectively. DNA extractions from fins were performed as in (Schulte et al., 2011). To extract DNA from fixed embryos, yolk was removed and heads dissected at the hindbrain border in 70% glycerol/ 30% PBS with insect pins. Trunks were stored in 70% glycerol/30% PBS at 4⁰C for analysis. Heads were incubated in 50 μL of Proteinase K solution for 2 hrs at 55°C. Proteinase K was heat inactivated at 100°C for 10 minutes and tubes were centrifuged for 20 minutes at 13000 rpm. DNA was precipitated with 100% ice-cold ethanol at −20°C overnight and re-suspended in 20 μL of water. Alternatively, DNA was extracted from dissected heads of fixed embryos using HotSHOT method (Truett et al., 2000), adding 20 μL of 50 mM NaOH and 2 μL of 1M Tris-HCl (pH 7.5). From the resuspended or extracted DNA, 2 μL was used for each PCR.

The *lbx1a^hu3569^* mutation creates a *XbaI* recognition site. Therefore, to genotype *lbx1a^hu3569^* mutants, a genomic region flanking the mutation was PCR-amplified using: 94°C for 60 seconds, followed by 5 cycles of 94°C 30 seconds, 54°C 30 seconds, 72°C 60 seconds; followed by 40 cycles of 92°C 20 seconds, 52°C 30 seconds, 72°C 60 seconds, and then a final extension at 72°C for 5 minutes. Forward primer: TTTACAGGCCTCTGCTGTTC. Reverse primer: AACACTCTTTGCTCGTTGTG. PCR products were digested with *XbaI* and analyzed on 1% TAE agarose gels. WT product is 510 bp. Mutant amplicons are cut into 256 bp and 254 bp fragments, usually detected as one band on the gel.

The *lbxlb^hu3534^* mutation creates a *AccI* recognition site. To genotype *lbx1a^hu3534^* mutants, a genomic region flanking the mutation was PCR-amplified with the same conditions as above using Forward primer: GCTATAGACAAAGGCTGGAATG and Reverse primer: GCCTACAATATACCCAGAATTG. PCR products were digested with *AccI* and analysed on 1% TAE agarose gels. WT product is 477 bp. Mutant amplicons are cut into 233 bp and 214 bp fragments. Alternatively, we used KASP assays (Biosearch Technologies). These use allele-specific PCR primers, which differentially bind fluorescent dyes, quantified with a Bio-Rad CFX96 real-time PCR machine to distinguish genotypes. Proprietary primer used was: *lbx1b_hu3534*.

To genotype *lbx1a^sa1496^* mutants, a base pair change adjacent to the mutation was introduced in the forward PCR primer to create a *ScaI* recognition site only in mutant DNA. The region flanking the mutation was PCR-amplified using: 94°C for 60 seconds, followed by 35 cycles of 94°C 30 seconds, 60°C 30 seconds, 72°C 45 seconds, followed by final extension at 72°C for 5 minutes. Forward primer: GAGAAAGTCGAGAACAGCTTTCACCAAGTAC. Reverse primer: CCTTCATCTCCTCTAGGTCTCTTTTGAGTT. PCR products were digested with *ScaI* and analysed on 2.5 % TBE agarose gels. WT product is 191 bp. Mutant amplicons are cut into 162 bp and 29 bp fragments. Alternatively, we used a KASP assay with proprietary primer *lbx1a_sa1496*.

### *in situ* hybridization on *Danio rerio*

Embryos were fixed in 4 % paraformaldehyde (PFA) and *in situ* hybridizations were performed as in (Batista et al., 2008; Concordet et al., 1996). Embryos older than 24 h were usually incubated in 0.003% 1-phenyl-2-thiourea (PTU) to prevent pigment formation. RNA probes were prepared using the following templates, *lbx1a* and *lbx2* (Ochi & Westerfield, 2009), *lbx1b* (Thisse et al., 2004), *dbx2* (Gribble et al., 2007), *pax2a* (Pfeffer et al., 1998).

To determine neurotransmitter phenotypes, we used *slc32a1* (formerly called *viaat*), which encodes a vesicular inhibitory amino acid transporter and labels all inhibitory cells (Kimura et al., 2006), a mixture of two probes (*glyt2a* and *glyt2b*) for *slc6a5* (previously called *glyt2*), which encodes a glycine transporter necessary for glycine reuptake and transport across the plasma membrane and labels glycinergic inhibitory cells (Higashijima et al., 2004) and a mixture of two probes to *gad1b* (previously called *gad67*) and one probe to *gad2* (previously called *gad65*) which label GABAergic inhibitory cells (Higashijima et al., 2004). *gad1b* and *gad2* encode for glutamic acid decarboxylases, necessary for the synthesis of GABA from glutamate. Glutamatergic (excitatory) cells were labelled with a mixture of *slc17a6b* (formerly called *vglut2.1*) and *slc17a6a* (formerly called *vglut2.2*; (Higashijima et al., 2004). These genes encode proteins responsible for transporting glutamate to the synapse. In all of these cases, a mix of equal concentrations of relevant probes was used.

### *in situ* hybridization on *Scyliorhinus canicula*

<u>*Scyliorhinus canicula*</u> (*S. canicula*) egg cases were obtained from Marine Biological Association, United Kingdom in Plymouth. Fertilized eggs were stored at 17°C in a 10 L aerated sea water container and staged according to (Ballard et al., 1993). The anterior and posterior tendrils from each egg case were cut and embryo position was determined by shining a bright light behind the egg case. A large window was cut where the embryo was located. The yolk stalk was pulled out using a pair of tweezers and cut with dissection scissors. The embryo was spooned out, washed with PBS, then placed in PBS with tricaine methanesulfonate (Sigma-Aldrich, A5040) until still, followed by fixation in 4% PFA and 3 washes for 5 minutes in PBS.

Cryosections were prepared by incubating fixed embryos in 30% sucrose in PBS at 4°C overnight. Embryos were trimmed and set into OCT on dry ice and then sectioned or stored at −20°C. Sections were cut on a Leica Jung Frigocut 2800E cryostat at approximately 20-40 μm thickness and collected on SuperFrost® Plus (Menzel-Gläser) slides and stored at −20°C. The zebrafish *in situ* protocol was used with the following modifications: slides were rehydrated in PBS or PBT, 200 μL of RNA probe in hybridization buffer was immediately placed onto sections and a coverslip was added, slides were incubated at 70°C in a sealed box overnight. Slides were placed in Coplin jars and washed as in zebrafish protocol but with the first formamide washes omitted. For staining, 500 μL of NBT/BCIP solution diluted in NTMT (0.1M Tris pH 9.5, 50mM MgCl2, 0.1M NaCl, 0.1% Tween 20) was placed on sections, and coverslipped slides were placed in the dark until staining developed. Then slides were washed in NTMT and PBS and fixed with 4% PFA.

*S. canicula lbx1* and *lbx2* correspond to cDNA fragments sequenced by Sanger sequencing. They were generated as part of a large-scale EST sequencing project of an *S. canicula* embryonic cDNA library (stages 9-15) as described in (Coolen et al., 2007). Lbx1 and Lbx2 sequences have been deposited in GenBank, with accession numbers MW456671 and MW456672 respectively. Recombinant plasmids were cut with SalI (*Lbx1*) and Kpn1 (*Lbx2*) and used to generate antisense RNA probes.

### *in situ* hybridization on *Xenopus tropicalis*

*Xenopus tropicalis* embryos were obtained from Jim Smith’s Lab at the University of Cambridge. Embryos were incubated at 25°C until the appropriate stage, when the vitelline membrane was removed by forceps, and the embryos were fixed in 4% PFA at 4°C overnight. Embryos were then washed in PBT and dehydrated in 100% methanol and stored at −20°C. Embryos in methanol were transferred to 100% ethanol and rehydrated through an ethanol/PBT series (90%, 70%, 50%, 30%, 10%), 5 minutes each. Rehydrated embryos older than stage 25 were incubated in 5 μg/ml proteinase K at room temperature for 15 minutes, followed by re-fixation in 4% PFA. Fixed embryos were placed into hybridization buffer (50% formamide, 5×SSC, 1 mg/ml yeast tRNA (Roche, 10109223001), 100 μg/ml heparin (Sigma-Aldrich, H9399), 2% Blocking reagent (Roche, 11096176001), 0.1% Tween 20, 0.1% CHAPS (Sigma-Aldrich)) until embryos sank and then incubated in fresh hybridization buffer for 5 hours at 70°C. This was followed by overnight incubation at 70°C with RNA probe (Martin & Harland, 2006) in hybridization buffer plus 0.1% SDS to enable probe penetration.

Embryos were washed in hybridization buffer at 70°C for 10 minutes, followed by three 20-minute washes in 2×SSC, 0.3% CHAPS at 60°C, two 30-minute washes in 0.2×SSC, 0.3% CHAPS and two 10 minute 0.3% CHAPS in PBT washes at 60°C. After a 10-minute wash in PBT at room temperature, embryos were incubated with 0.5% blocking reagent in PBT before an overnight incubation in 1:2000 anti-dig AP antibody (Roche, 11093274910) in 0.5% blocking reagent (Roche, 11096176001), in PBT at 4°C.

After antibody incubation, embryos were washed five times for one hour, in PBT. Then, they were transferred into a 24-well plate and washed twice in NTMT for five minutes. Color reaction was performed by adding 20 μl/ml NBT/BCIP per ml of NTMT and placing embryos in the dark. To stop staining, embryos were washed several times in PBT and fixed in 4% PFA. When required, pigment was removed by washing embryos four times in 70% ethanol in PBS for one hour and then placing in bleach (3% H2O2, 5% formamide, 0.5×SSC) for 5 minutes, followed by incubating for 2 hours on a light box with fresh bleach and then washing several times with PBS. Prior to staining visualization, embryos were dehydrated in several washes of methanol and transferred into glass watch glasses where they were cleared in Murray’s solution (2:1 benzyl benzoate : benzyl alcohol) (Klymkowsky & Hanken, 1991).

### *in situ* hybridization plus imunohistochemistry on *Danio rerio*

Primary antibodies used were chicken polyclonal anti-GFP (Abcam, ab13970, 1:500), rabbit anti-GFP (Molecular Probes A6465, 1:500) and rabbit anti-activated Caspase-3 (Fisher Scientific/BD, BDB559565, 1:500). Antibody used for fluorescent *in situ* hybridization was mouse anti-Dig (Jackson ImmunoResearch 200-002-156, 1:5000), detected with Invitrogen Tyramide #5 (ThermoFisher Scientific, T20915). Secondary antibodies used were Alexa Fluor 568 goat anti-mouse (ThermoFisher Scientific, A-11031, 1:500), Alexa Fluor 488 goat anti-rabbit (ThermoFisher Scientific, A-11034, 1:500) and Alexa Fluor 488 goat anti-chicken IgG (H+L) (ThermoFisher Scientific, A-11039, 1:500).

Embryos for immunohistochemistry were treated with acetone for 20 min to permeabilize them, then washed for 5 min in distilled water and 2 x 10 min in PBS. Embryos were treated with Image-iT Signal Enhancer (ThermoFisher Scientific, I36933) for 30 min, then incubated in block solution (2 % goat serum, 1 % BSA, 10 % DMSO and 0.5 % Triton) for 1 h followed by incubation in primary antibody in fresh block solution at 4°C overnight. Embryos were washed with PBT (PBS + 0.1 % Triton) for 2 h and incubated with secondary antibody in block solution at 4°C overnight. Embryos were then washed with PBT for 2 h and stored in 2 % DABCO (Acros Organics, AC112471000).

### Image acquisition and processing

Whole-mount tadpoles were placed in a 1% agarose plate and covered in PBS for imaging using an Olympus SZX16 stereomicroscope and a Q-Imaging Micropublisher 5.0 RTV camera. Whole mount zebrafish embryos and *S. canicula* cross-sections were mounted in either 70% glycerol, Vectashield or 2% DABCO on a microscope slide. DIC pictures were taken using an AxioCam MRc5 camera mounted on a Zeiss Axio Imager M1 compound microscope. Fluorescence-only images were taken on a Zeiss LSM 710 confocal microscope. Images were processed using Adobe Photoshop software (Adobe, Inc) and Image J software (Abràmoff et al., 2004). Combined fluorescent and brightfield images were merged in Photoshop by placing fluorescent images on top of brightfield images and adjusting opacity and/or fill of the fluorescent image.

### Cell counting and statistical analyses

In all cases, cells counts are for both sides of a five-somite length of spinal cord adjacent to somites 6-10. Data were analyzed for normality using Shapiro-Wilk test in R version 3.5.1 (R_Development_Core_Team, 2005). All data sets analyzed had normal distributions. For pairwise comparison of *slc17a6* expression in WT and *lbx1a^hu3569^* mutant embryos, the F-test for equal variances was performed, and as variances were equal, a type 2 (for equal variances) student’s *t*-test was performed. To control for type I errors in all other data sets comparing WT, *lbx1a*, *lbx1b* and *lbx1a;lbx1b* mutant embryos, a one-way analysis of variance (ANOVA) test was performed. Data sets were first assessed for homogeneity of variances using Bartlett’s test. All had homogeneous (homoscedastic, Bartlett’s test p value >0.05) variances and so standard ANOVA analysis was performed. ANOVA results are reported as F(dfn,dfd) = f-ratio, p value = *x*, where F = F-statistic, dfn = degree of freedom for the numerator of the F-ratio, dfd = degree of freedom for the denominator of the R-ratio, and *x* = the p value. For statistically significant ANOVA, to determine which specific groups differed, Tukey’s honestly significant difference post hoc test for multiple comparisons was performed. F-test, and student’s t-test were performed in Microsoft Excel version 16.41. Bartlett’s testing, standard ANOVA, and Tukey’s honestly significant difference testing were performed in Prism version 9.0.0 (GraphPad Software, San Diego, California USA, www.graphpad.com).

## Results

### Zebrafish *lbx* genes have different spinal cord expression patterns

In contrast to amniotes which have two *Lbx* genes, *Lbx1* and *Lbx2*, teleosts have two *lbx1* genes, *lbx1a* and *lbx1b*, and one *lbx2* gene (Wotton et al., 2008; Wotton et al., 2010). In addition, while only *Lbx1* is expressed in amniote spinal cord, all three zebrafish *lbx* genes are expressed in spinal cord (Fig. 2; Lukowski et al., 2011; Neyt et al., 2000; Ochi & Westerfield, 2009). To examine and compare spinal expression of the three zebrafish *lbx* genes we performed *in situ* hybridizations. Our data show that *lbx1a*, *lbx1b* and *lbx2* are all expressed in zebrafish spinal cord by mid-somitogenesis stages. At 18h (18-somites) *lbx1a* is expressed in a subset of dorsal spinal cord cells. This expression is stronger rostrally, and decreases more caudally (Fig. 2a). As development proceeds, additional cells in the same dorsal domain start to express *lbx1a* and expression extends more caudally (Fig. 2b & d). By 30h, *lbx1a* is expressed along the whole rostral-caudal axis of the spinal cord (Fig. 2d). Analyses of spinal cross-sections show that *lbx1a*-expressing cells are located at the lateral edges of dorsal spinal cord, consistent with expression by post-mitotic interneurons (Fig. 2c; at these stages of development, progenitor cells are located medially next to the ventricle and post-mitotic neurons are located at the lateral edge of the spinal cord). Further confirming that this expression is in post-mitotic interneurons, *lbx1a* spinal expression is expanded in *mindbomb1^ta52b^* mutants at 24h (Fig. 2e & f). *mindbomb1* is a ubiquitin-ligase essential for efficient Notch signaling. When Notch signaling is disrupted or lost, spinal progenitor cells precociously differentiate as early forming neurons, resulting in a loss of progenitor gene expression and expanded expression of most post-mitotically-expressed genes (e.g. Batista et al., 2008; Jiang et al., 1996; Park & Appel, 2003).

**Figure 2.**
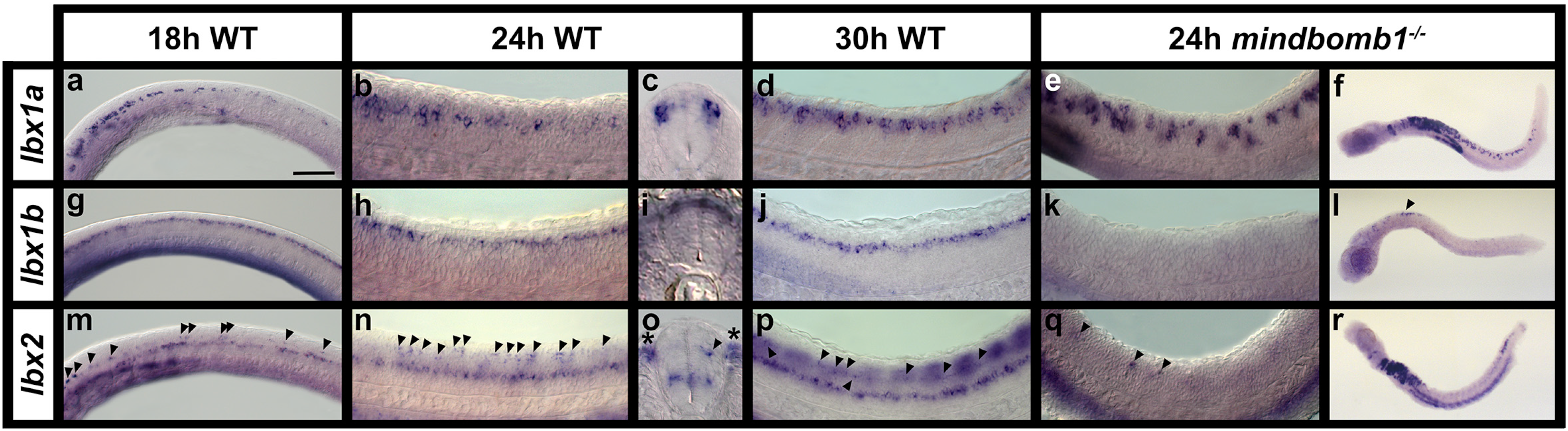
Expression of *lbx* genes in zebrafish spinal cord. **(a, b, d, e, g, h, j, k, m, n, p & q)** Lateral views of spinal cord expression of *lbx* genes at 18h (18-somites; **a, g, m)**, 24h **(b, e, h, k, n** & **q)** and 30h **(d, j, p)** in WT embryos **(a-b, d, g-h, j,** & **m-n & p)** and *mindbomb1^ta52b^* mutants **(e, k** & **q)** and lateral views of whole *mindbomb1^ta52b^* mutants at 24h (**f, l & r**). Rostral is left and dorsal up in all cases. **(c, i & o)** Spinal cord cross-sections of 24h WT embryos. **(a)** *lbx1a* is expressed in the hindbrain and rostral spinal cord at 18h, and caudally in a few scattered dorsal spinal cord cells. At 24h **(b)** and 30h **(d)**, expression extends more caudally. **(c)** Cross-section of WT spinal cord confirms that *lbx1a*-expressing cells are located laterally in a post-mitotic dorsal spinal cord domain. **(e)** *lbx1a* spinal expression is expanded in *mindbomb1^ta52b^* mutant embryos at 24h, again suggesting that the cells expressing this gene are post-mitotic. **(f)** At 24h, this expanded expression is most pronounced in the rostral spinal cord. **(g)** *lbx1b* is expressed in an almost continuous row of cells in the hindbrain and dorsal spinal cord at 18h. This expression persists at 24h and 30h **(h** & **j)**. **(i)** Cross-section of WT spinal cord shows *lbx1b* expression both medially and laterally in dorsal spinal cord, suggesting it is expressed by both post-mitotic and progenitor cells. **(k)** *lbx1b* expression is lost throughout most of the spinal cord in *mindbomb1^ta52b^* mutants, suggesting that this gene is expressed by progenitor cells that differentiate precociously in these mutants. **(l)** A small number of cells still express *lbx1b* in the very rostral spinal cord (black arrow head). It is unclear why this region differs from the rest of the spinal cord. **(m)** At 18h, *lbx2* is expressed in a continuous row of cells in ventral spinal cord and discontinuously in a more dorsal row of cells (indicated with black arrowheads, which point to some of the expressing cells in this dorsal row). **(n)** This expression remains at 24h. **(o)** Cross-sections of WT spinal cord at this stage confirm that the ventral *lbx2*-expressing spinal cord cells are mainly located medially (suggesting they are likely progenitor cells), whereas dorsal *lbx2*-expressing cells (black arrow head) are more lateral (suggesting they are either becoming, or are already, post-mitotic cells). Some are (like in **o**) located slightly medial to the lateral edge of the spinal cord and some are located at the lateral edge. Somite staining can also be observed outside of the spinal cord (indicated with black asterisks). **(p)** At 30h, expression of *lbx2* in the dorsal spinal cord becomes more difficult to see due to strong somite staining (seen here as out of focus repeated blocks over dorsal spinal cord). **(q)** *lbx2* expression is lost in the ventral spinal cord domain in *mindbomb1^ta52b^* mutants, although a small number of *lbx2*-expressing cells remain more dorsally. This suggests that if *lbx2* is expressed by any post-mitotic cells, then it is only for a short period of time. (**r**) There is also expanded expression of *lbx2* in the hindbrain and caudal dorsal spinal cord. The caudal expression is likely to be post-mitotic cells that have not yet turned *lbx2* expression off. Scale bar: 50 μm (**a, b, d, e, g, h, j, k, m, n, p & q**), 200 μm (**f, l & r**), 30 μm (**c, i, o**).

*lbx1b* is also expressed in dorsal spinal cord at 18h, but unlike *lbx1a*, it is expressed by a continuous row of cells along the rostral-caudal axis, similar to progenitor domain genes, and its expression extends more caudally than *lbx1a* (Fig. 2g). This expression pattern persists at 24 and 30h (Fig. 2h & j). Analyses of spinal cross-sections suggest that *lbx1b* is expressed in both medial progenitor and lateral post-mitotic spinal cells. This suggests that *lbx1b* expression may persist, at least for a short time, in post-mitotic interneurons (Fig. 2i). However, in 24h *mindbomb1^ta52b^* mutants, most of the spinal expression of *lbx1b* is lost, consistent with it being expressed in progenitor cells and suggesting that if it is expressed in post-mitotic cells, it is very quickly turned off after these cells become post-mitotic (Fig. 2k & l).

In contrast to *lbx1a* and *lbx1b*, *lbx2* is expressed in two different dorso-ventral spinal domains. At 18h, the dorsal row of *lbx2* expression consists of fewer, more spaced cells than the more continuous ventral row (Fig. 2m) and this expression pattern persists at 24h and 30h (Fig. 2n & p). The dorsal row is most clearly visible in the rostral spinal cord. *lbx2* is also expressed in rostral somites at 18h and this expression extends caudally and increases by 24h and 30h, making it harder to clearly see spinal expression (Fig. 2m-p). Analyses of spinal cross-sections at 24h show that ventral *lbx2*-expressing spinal cells are predominantly medial, although there are also occasional lateral cells, and dorsal *lbx2*-expressing cells are located either at the lateral edges of the spinal cord or between the medial ventricle and the lateral edge of the spinal cord (Fig. 2o). This suggests that the dorsal *lbx2* expression domain consists of cells that are becoming post-mitotic and the ventral expression domain is predominantly progenitor cells. In *mindbomb1^ta52b^* mutants at 24h, most *lbx2* spinal expression is lost (Fig. 2 q & r), although there is an expansion in the number of cells expressing *lbx2* in the caudal spinal cord (Fig. 2r). This is consistent with ventral *lbx2*-expressing cells being predominantly progenitor cells and it suggests that even in the more dorsal domain of expression, *lbx2* expression is turned off soon after cells become post-mitotic.

Unfortunately, the zebrafish *lbx in situ* probes are relatively weak and, as a result, we were unable to successfully perform double *in situ* hybridizations with combinations of these genes. Therefore, to compare different *lbx* spinal expression domains, we identified a putative 1.6 Kb enhancer region (CNE) downstream of *lbx1a* and constructed the *Tg(1.6 lbx1a:EGFP)^SU33^* transgenic line (see materials and methods & Fig. 1). This line recapitulates endogenous *lbx1a* expression (Fig. 3a & b). EGFP is expressed in the same dorso-ventral spinal region as endogenous *lbx1a* mRNA and at least most of the EGFP-positive cells co-express *lbx1a* mRNA. In contrast, a different transgenic line, *Tg(0.9 lbx1a:EGFP)^SU32^*, that we constructed using a smaller 900bp putative enhancer region that is located closer to the 3’ end of *lbx1a* (see materials and methods & Fig. 1), was expressed in relatively few spinal cord cells (Fig. 3g). We also confirmed that a previously published *Tg(lbx1b:EGFP)^ua1001^* line (Lukowski et al., 2011) is expressed in a similar dorsal spinal domain to endogenous *lbx1b* expression (Fig. 3j; Lukowski and colleagues reported that this line recapitulates endogenous *lbx1b* expression but did not show supporting data).

**Figure 3.**
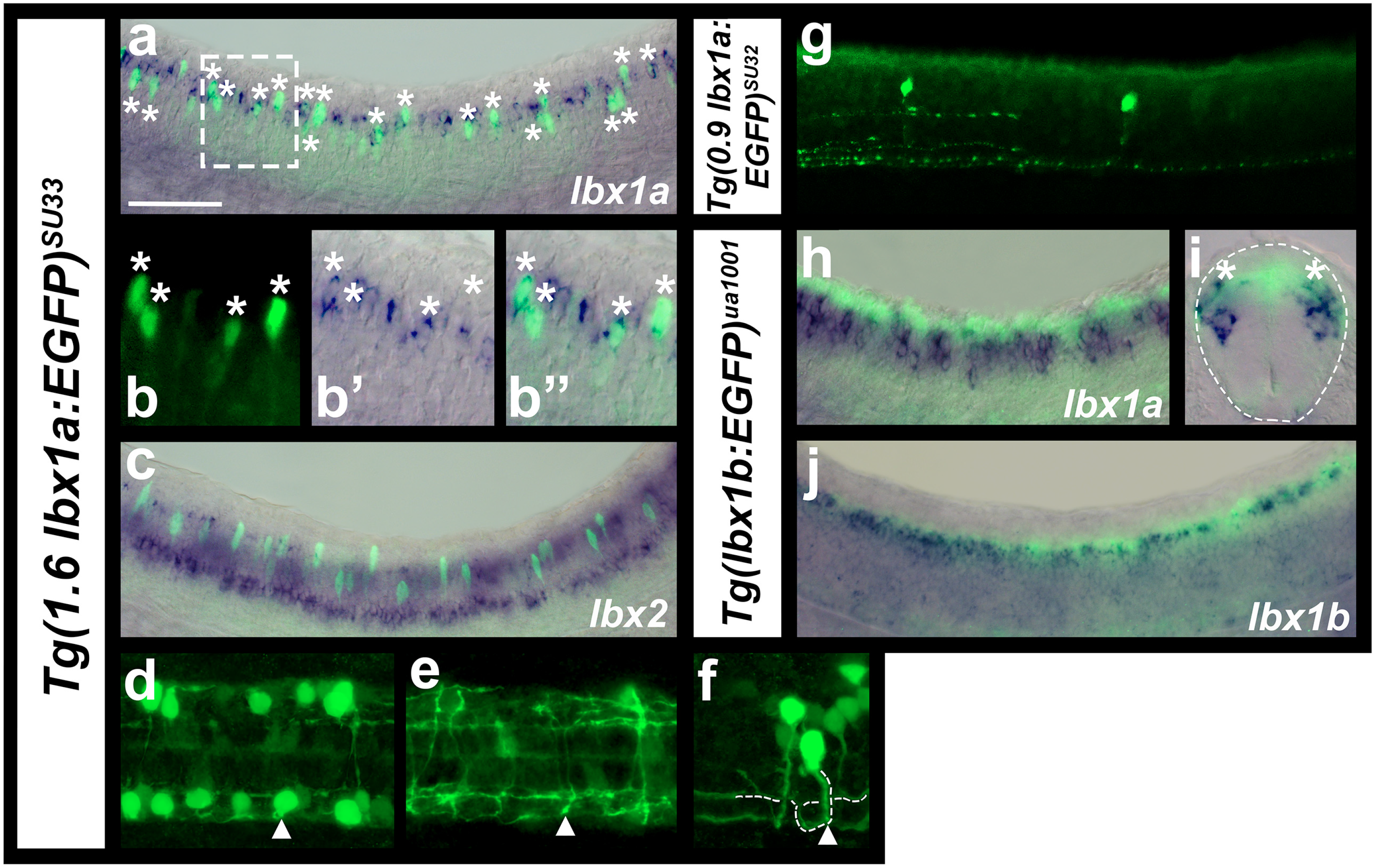
Comparisons of zebrafish *lbx1a*, *lbx1b* and *lbx2* spinal cord expression. Immunohistochemistry for EGFP (green) in *Tg(1.6 lbx1a:EGFP)^SU33^* (**a, b & f-i-f**), *Tg(0.9 lbx1a:EGFP)^SU32^*(**g**) and *Tg(lbx1b:EGFP)^ua1001^* (**h-jc-e**) embryos, coupled with *in situ* hybridization (blue) for *lbx1a* (**a, b, h-c & ie**), *lbx1b* (**gd**) or *lbx2* (**cg**). Lateral views of spinal cord with dorsal up and anterior left at 27h (**h, jc-d**), 30h (**a-c, b, g**) and 35h (**f, g**), cross-section of spinal cord at 27h (**ie**) and dorsal views of two different focal planes of the spinal cord at 35h (**dh & ei**). White dotted box in (**a**) indicates the region shown in magnified view of a single confocal plane in (**b**). White asterisks in (**a & b**) indicate co-labeled cells. Occasional single-labeled EGFP cells may be the result of weak endogenous *lbx1a* mRNA expression not being detected in the double staining experiment or they may be cells that used to express *lbx1a* and the EGFP expression has persisted. Single-labeled *lbx1a* mRNA-expressing cells are probably cells that have turned on *lbx1a* expression but not yet made EGFP protein. It is also possible that the NBT/BCIP precipitate has quenched the fluorescent signal in these cells. In contrast, we did not observe any co-labeled *lbx2* and *Tg(1.6 lbx1a:EGFP)^SU33^* cells (**c**). (**d-f**) white arrowhead (**d**) indicates the same neuron whose axon goes ventral in the spinal cord (**e**), crosses the midline and bifurcates on the other side of the spinal cord (**f**). (**f**) Dotted white line (drawn slightly to the right of the axon so EGFP expression is still visible) indicates commissural bifurcating axon trajectory. White arrowhead indicates where the axon starts to cross the midline. (**g**) The shorter 0.9 Kb *lbx1* CNE (Figure 1), used to make the *Tg(0.9 lbx1a:EGFP)^SU32^* transgenic line, only drives *lbx1* expression in very few spinal cord neurons. Dotted line (**ie**) indicates edge of the spinal cord. Co-expression of *Tg(lbx1b:EGFP)^ua1001^* and *lbx1a* can be seen in the dorsal-most region of the lbx1a-expression domain (white asterisks in **ie**). (**j**) *Tg(lbx1b:EGFP)^ua1001^* is co-expressed in a same dorsal spinal domain as endogenous *lbx1b*. Scale bar: 50 μm (**a, c, g, hd & jg**), 35 μm (**ie**), 25 μm (**b**, **df, eh & fi**).

When we compare expression of *lbx1a* and *lbx1b* to *Tg(lbx1b:EGFP)^ua1001^* it is clear that *lbx1a* spinal expression is, in the main, more ventral than that of *lbx1b* although the two genes overlap in the most dorsal region of the *lbx1a* expression domain (Fig. 3h-j). Comparisons of *lbx1a* and *lbx2* to *Tg(1.6 lbx1a:EGFP)^SU33^* also confirm that the ventral row of *lbx2* expression is more ventral than *lbx1a* expression although the dorsal *lbx2*-expressing cells are located at a similar dorsal-ventral position to some of the *lbx1a*-expressing cells (Fig. 3c).

### Zebrafish *lbx1a*-expressing cells develop into commissural bifurcating interneurons (CoB)

Previous work in mouse has shown that *Lbx1* is expressed by dI4, dI5 and dI6 interneurons and subsequently by later forming dILA and dILB interneurons (Gross et al., 2002; Müller et al., 2002). However, the axon trajectories and morphologies of these cells have not been described in detail, although data from Gross and colleagues suggest that many of the later-born cells are ipsilateral (Gross et al., 2002). When we examined *Tg(1.6 lbx1a:EGFP)^SU33^* embryos we found that by 35 h, at least most of the EGFP-positive cells have extended their axons ventrally and crossed the midline to the other side of the spinal cord (Fig. 3 d-f). We determined the axon trajectories of 66 GFP-positive spinal neurons and found that all of these turned slightly dorsally and then bifurcated after they crossed the midline. This suggests that many *lbx1a*-expressing spinal interneurons have a commissural bifurcating, or CoB morphology (Fig. 3 d-f).

### Zebrafish *lbx1a* is expressed by dI4, dI5 and dI6 spinal interneurons

As zebrafish *lbx1a* seemed to be expressed in a similar spinal domain to mouse *Lbx1*, we used double-labeling experiments to test whether it is expressed by dI4, dI5 and dI6 interneurons. dI4, dI5 and dI6 spinal interneurons are located immediately dorsal to V0 interneurons and develop both from, and dorsal to, the *dbx2*-expressing progenitor domain (Lewis, 2006 and references therein). We found that some *Tg(1.6 lbx1a:EGFP)^SU33^*-expressing cells are at the same dorso-ventral level as *dbx2*-expressing cells and some are more dorsal, although, as expected for post-mitotic interneurons, the EGFP-expressing cells are lateral to the *dbx2*-expressing cells (Fig. 4a & b). In addition, when we compared the expression of *lbx1a* to *Tg(evx1:EGFP)^SU1^*, which labels V0v interneurons (Juárez-Morales et al., 2016), we found that most of the *lbx1a*-expressing cells are dorsal to V0v interneurons and we did not observe any co-expression of *lbx1a* and EGFP (Fig. 4c & d). We also compared expression of *Tg(1.6 lbx1a:EGFP)^SU33^* to *pax2a*, which is expressed by V1, V0D, dI4 and dI6 spinal interneurons (Batista & Lewis, 2008). We found that most of the *lbx1a*-expressing cells are located in the same dorso-ventral spinal region as *pax2a*-expressing cells, and more importantly, a subset of EGFP-positive cells co-express *pax2a* (Fig. 4e). Finally, as dI4 and dI6 interneurons are inhibitory and dI5 interneurons are excitatory (Cheng et al., 2005; Gross et al., 2002; Müller et al., 2002), we also examined neurotransmitter phenotypes of *Tg(1.6 lbx1a:EGFP)^SU33^*-expressing cells. We found that many *Tg(1.6 lbx1a:EGFP)^SU33^*-expressing cells co-express the inhibitory marker *slc32a1* (*viaat;* Fig. 4f), and a smaller number co-express the excitatory marker *slc17a6* (*vglut2;* Fig. 4g; see materials and methods for a more detailed discussion of neurotransmitter markers used). Taken together, these data suggest that, like mouse *Lbx1*, zebrafish *lbx1a* is expressed in dI4, dI5 and dI6 spinal interneurons.

**Figure 4.**
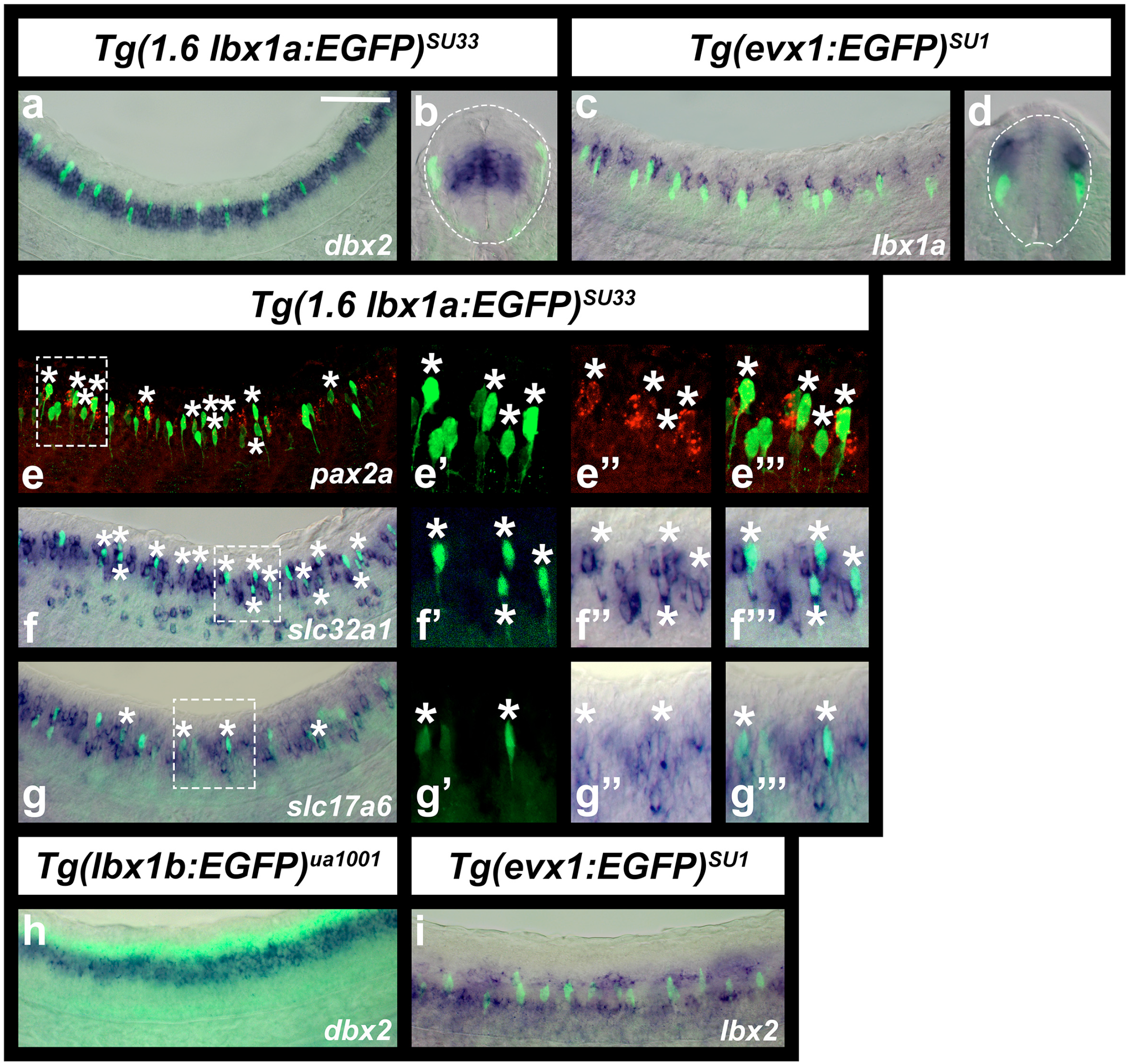
Zebrafish *lbx1a* is expressed by dI4, dI5 and dI6 spinal interneurons. **(a-i)** Immunohistochemistry for EGFP (green) in *Tg(1.6 lbx1a:EGFP)^SU33^* (**a, b & e-g**), *Tg(evx1:EGFP)^SU1^* (**c, d & i**), and *Tg(lbx1b:EGFP)^ua1001^* (**h**) embryos, coupled with *in situ* hybridization (blue) for *dbx2* **(a, b** & **h)**, *lbx1a* **(c** & **d)**, *slc32a1* **(f)**, *slc17a6* **(g)**, *lbx2* **(i)**, and *in situ* hybridization (red) for *pax2a* **(e)**. *dbx2* **(a, b** & **h)** is expressed in dP6, p0 and p1 progenitor domains, whereas *pax2a* **(e)** is expressed by V1, V0D, dI4 and dI6 spinal interneurons and *evx1* **(c, d & i** is expressed by V0v spinal inteneurons. Lateral views with dorsal up and anterior left of spinal cord at 30h **(a & e-h)** and 24h **(c & i)** and cross-sections with dorsal up at 30h (**b**) or 24h (**d**). (**e-g)** panels on the right are magnified views of single confocal planes of white dotted box region in left-hand panel. White asterisks indicate co-labeled cells. **(b** & **d)** White dotted lines indicate the edge of the spinal cord. Scale bar: 50 μm (**a, c & e-i**), 35 μm (**b, d**, **e’, e’’, e’’’, f’, f’’, f’’’, g’, g’’, g’’’**).

Consistent with our comparisons of *lbx1a*, *lbx1b* and *lbx2* expression discussed above, *Tg(lbx1b:EGFP)^ua1001^* is expressed immediately dorsal to *dbx2* (Fig. 4h), and the ventral row of *lbx2*-expressing cells is located ventral to V0v interneurons and the dorsal row of *lbx2*-expressing cells is dorsal to these cells. (Fig. 4i). We did not observe any co-expression of *Tg(evx1:EGFP)^SU1^* and *lbx2.* These data suggest that *lbx1b* is probably expressed in the dP4 progenitor domain and the ventral *lbx2*-expressing cells are probably in the p1 progenitor domain.

### Lbx1a and Lbx1b are required to specify correct neurotransmitter fates of a subset of dorsal spinal interneurons

In mouse *Lbx1* mutants there is a reduction in the number of spinal GABAergic interneurons and a corresponding increase in spinal glutamatergic interneurons (Gross et al., 2002; Müller et al., 2002). To test whether this function of Lbx1 is conserved in zebrafish, we analyzed neurotransmitter phenotypes of *lbx1a* mutants. At 24h, we observed a slight reduction in the number of cells expressing *slc32a1* (previously called *viaat*), although this decrease was not statistically significant (Fig. 5m; Table 1). However, the reduction in the number of *slc32a1*-expressing cells became more pronounced and statistically significant at 30h (Fig. 5n; Table 1). In addition, there was a statistically significant increase in the number of spinal cells expressing *slc17a6* (previously called *vglut2*) at this stage (Fig. 5 g, h, o & p, Table 1).

**Figure 5.**
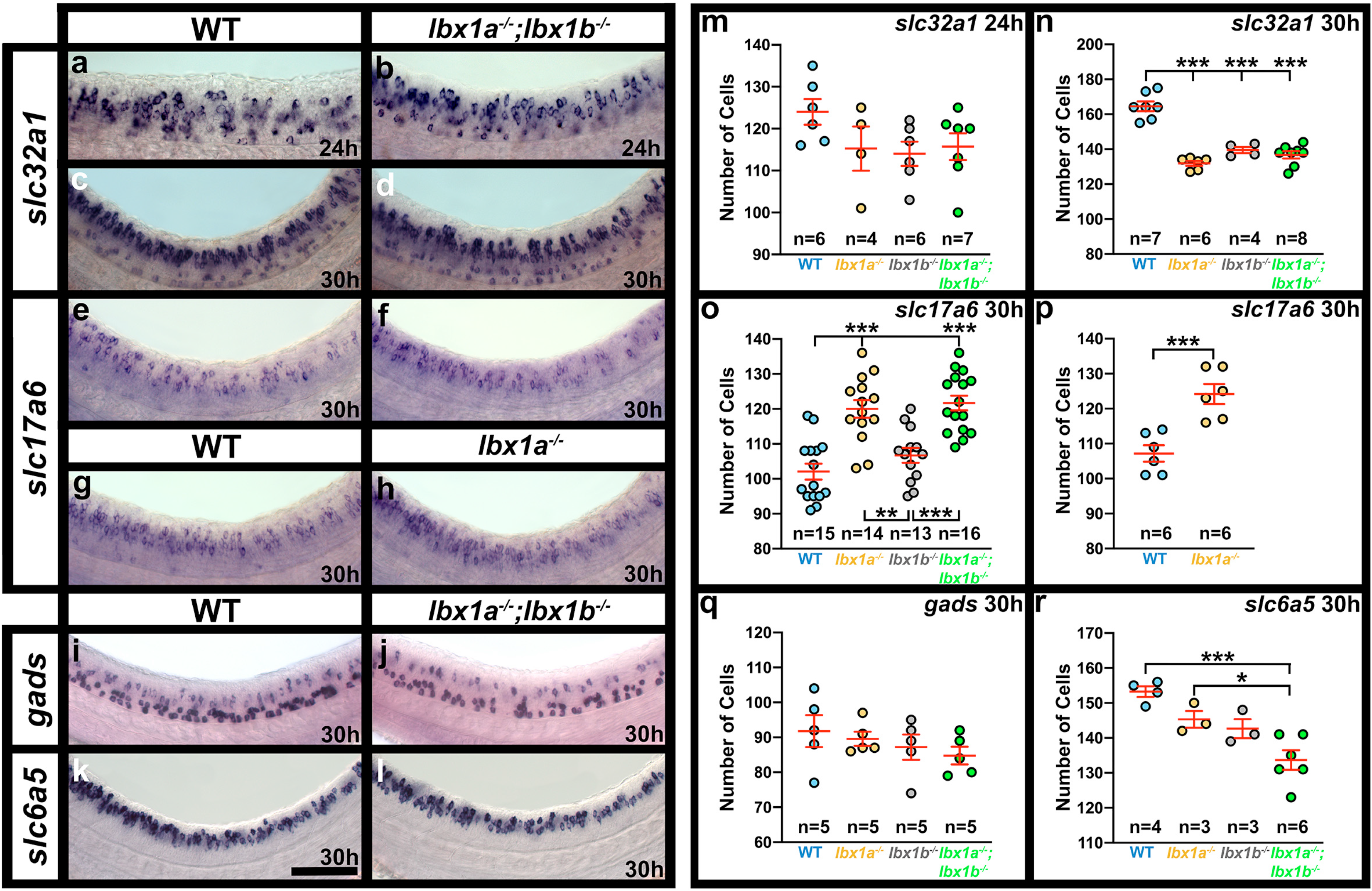
A subset of spinal interneurons have changed neurotransmitter phenotypes in the absence of Lbx1a and Lbx1b function. Expression of markers of different neurotransmitter phenotypes, *slc32a1* (also called *viaat*, marker of all inhibitory interneurons)*, slc17a6* (also called *vglut*, marker of all excitatory interneurons)*, gads* (marker of GABAergic inhibitory interneurons) and *slc6a5* (also called *glyt2*, marker of glycinergic inhibitory interneurons), in *lbx1a^−/−^, lbx1b^−/−^* single and double mutant embryos. Lateral views of zebrafish spinal cord at 24h **(a & b)** and 30h **(c-l),** showing *in situ* hybridization expression for genes indicated on the left. Anterior left, dorsal up. Mutant alleles are **(b)** *lbx1a^hu3569^;lbx1b^hu3534^*, **(d, f, j & l)** *lbx1a^sa1496^;lbx1b^hu3534^* and **(h)** *lbx1a^hu3569^*. *lbx1a^hu3569^* and *lbx1a^sa1496^* have similar phenotypes (compare **o & p**). **(m-r)** Number of cells (y-axis) expressing specific genes (indicated at top right) in spinal cord region adjacent to somites 6-10 at 24h **(m)** and 30h **(n-r)**. All data were first analyzed for normality using the Shapiro-Wilk test. All data sets are normally distributed. For the pairwise comparison shown in (**p**), the F test for equal variances was performed. This data set has equal variances and so a type 2 (for equal variances) student’s t-test was performed. To accurately compare the 4 different data sets shown in each of panels **m, n, o, q** and **r**, a one-way analysis of variance (ANOVA) test was performed. All data sets for ANOVA analysis have both normal distributions and homogeneous (homoscedastic, Bartlett’s test p value >0.05) variances and so standard ANOVA analysis was performed. The ANOVA results are as follows, only the ANOVA for panels **n, o** and **r** are significant (**m**: ANOVA (*F*(3,197) = 1.812, *p* = 0.1793), **n**: ANOVA (*F*(3,21) = 45.60, *p* = <0.0001), **o**: ANOVA (*F*(3,54) = 18.79, *p* = <0.0001), **q**: ANOVA (*F*(3,16) = 0.8174, *p* = 0.5030), **r**: ANOVA (*F*(3,12) = 11.05, *p* = 0.0009), and so to determine which specific experimental group or groups differed, Tukey’s honestly significant difference post hoc test for multiple comparisons was performed. Data are depicted as individual value plots and the *n*-values (number of embryos counted) are also indicated for each genotype. In each plot, the wider, middle red horizontal bar depicts the mean number of cells and the narrower red horizontal bars depict the standard error of the mean (S.E.M.). Statistically significant comparisons are indicated with brackets and asterisks. *p* < 0.05 = *, *p* < 0.001 = ***. Mean, S.E.M. and p values for comparisons are provided in Table 1. Scale bar: 50 μm **(a-l)**.

**Table 1.**
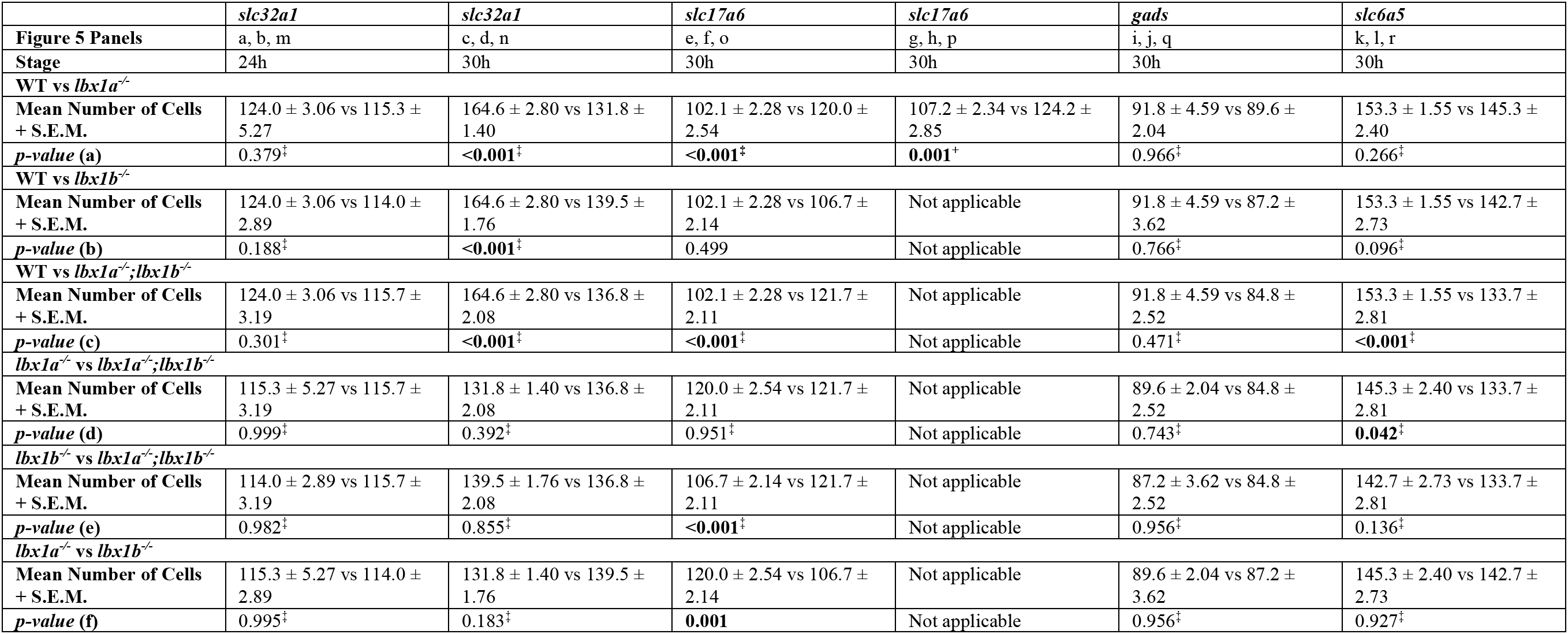
Number of cells expressing particular genes in WT and *lbx1* mutant embryos. Numbers of cells expressing particular genes (row 1) in spinal cord region adjacent to somites 6-10 and p values of comparisons between embryos with different genotypes (rows 6, 9, 12, 15, 18 and 21). Row 2 indicates the panels in Figure 5 that show the data for that comparison and row 3 indicates the developmental stage assayed. Rows 4, 7, 10, 13, 16 and 19 indicate the genotypes being compared. Values (rows 5, 8, 11, 14, 17 and 20) indicate the mean from at least 3 different embryos ± standard error of the mean (S.E.M.) for each genotype in that particular comparison. For all genes assayed in both *lbx1a/b* single and double mutants, a standard ANOVA was performed with Tukey’s honestly significant difference post-hoc testing. P values generated by this method of testing are indicated by (^‡^). For the 30h *slc17a6* experiment performed on *lbx1a^hu3569^* mutant embryos, a type 2 student’s t-test was performed, with p values indicated by (^+^). Statistically significant (p<0.05) values are indicated in bold. For a discussion of why particular tests were used, see materials and methods. P(a) (row 6) compares WT with *lbx1a* single mutant embryos, P(b) (row 9) compares WT with *lbx1b* single mutant embryos, P(c) (row 12) compares WT with *lbx1a;lbx1b* double mutant embryos, P(d) (row 15) compares *lbx1a* single mutant embryos with *lbx1a;lbx1b* double mutant embryos, P(e) (row 18) compares *lbx1b* single mutant embryos with *lbx1a;lbx1b* double mutant embryos and P(f) (row 21) compares *lbx1a* single mutant embryos with *lbx1b* single mutant embryos. Mean cell count values are provided to one decimal place, S.E.M. values to 2 decimal places and p values to three decimal places. The *lbx1a^sa1496^* allele was used in all experiments except the 24h *slc32a1* double mutant experiment and the 30h *slc17a6* single mutant experiment, in which the *lbx1a^hu3659^* allele was used instead. In all cases, the *lbx1b* allele used was *lbx1b^hu3534^*.

As the spinal expression patterns of *lbx1a* and *lbx1b* suggest that at least some *lbx1a*-expressing interneurons may develop from *lbx1b*-expressing progenitor cells, we also tested whether there was any redundancy between *lbx1a* and *lbx1b* by examining neurotransmitter phenotypes of *lbx1a;lbx1b* double mutants. There was no statistically significant difference between the number of spinal cells expressing either *slc32a1* or *slc17a6* in *lbx1a* single mutants compared to *lbx1a;lbx1b* double mutants (Fig. 5 f, h, n & o). However, while there was no increase in the number of spinal cells expressing *slc17a6* in *lbx1b* single mutants (Fig. 5o), there was a decrease in the number of spinal cord cells expressing *slc32a1* that was equivalent to that in *lbx1a* single mutants and *lbx1a;lbx1b* double mutants (Fig. 5n). This suggests that both *lbx1b* and *lbx1a* are required, presumably in succession, for the inhibitory fates of at least some dI4 and dI6 interneurons, but only *lbx1a* is required to repress excitatory fates in these cells.

To determine whether the reduction in the number of inhibitory cells in *lbx1a* and *lbx1b* single and double mutants represents a reduction in the number of GABAergic or glycinergic interneurons, we examined expression of markers of these neurotransmitter phenotypes in both single and double *lbx1a;lbx1b* mutants compared to WT embryos. There was no significant difference in the number of cells expressing GABAergic markers at 30h (Fig. 5j & q, Table 1). However, in contrast, there is a statistically significant decrease in the number of spinal interneurons expressing glycinergic markers in *lbx1a;lbx1b* double mutants (Fig. 5l & r, Table 1), although this reduction is less than the reduction in the number of cells expressing *slc32a1*.

To test whether the reduction in inhibitory interneurons might be caused by cell death, we performed activated caspase-3 immunohistochemistry. However, we did not observe any difference in the number of activated caspase-3 cells when comparing WT and double mutant embryos (*p* = 0.68; n= 3; Fig. 6). This is also consistent with the fact that there is an increase in the number of glutamatergic cells in the spinal cord of *lbx1a;lbx1b* double mutants, which suggests that cells are changing aspects of their fate rather than dying.

**Figure 6.**
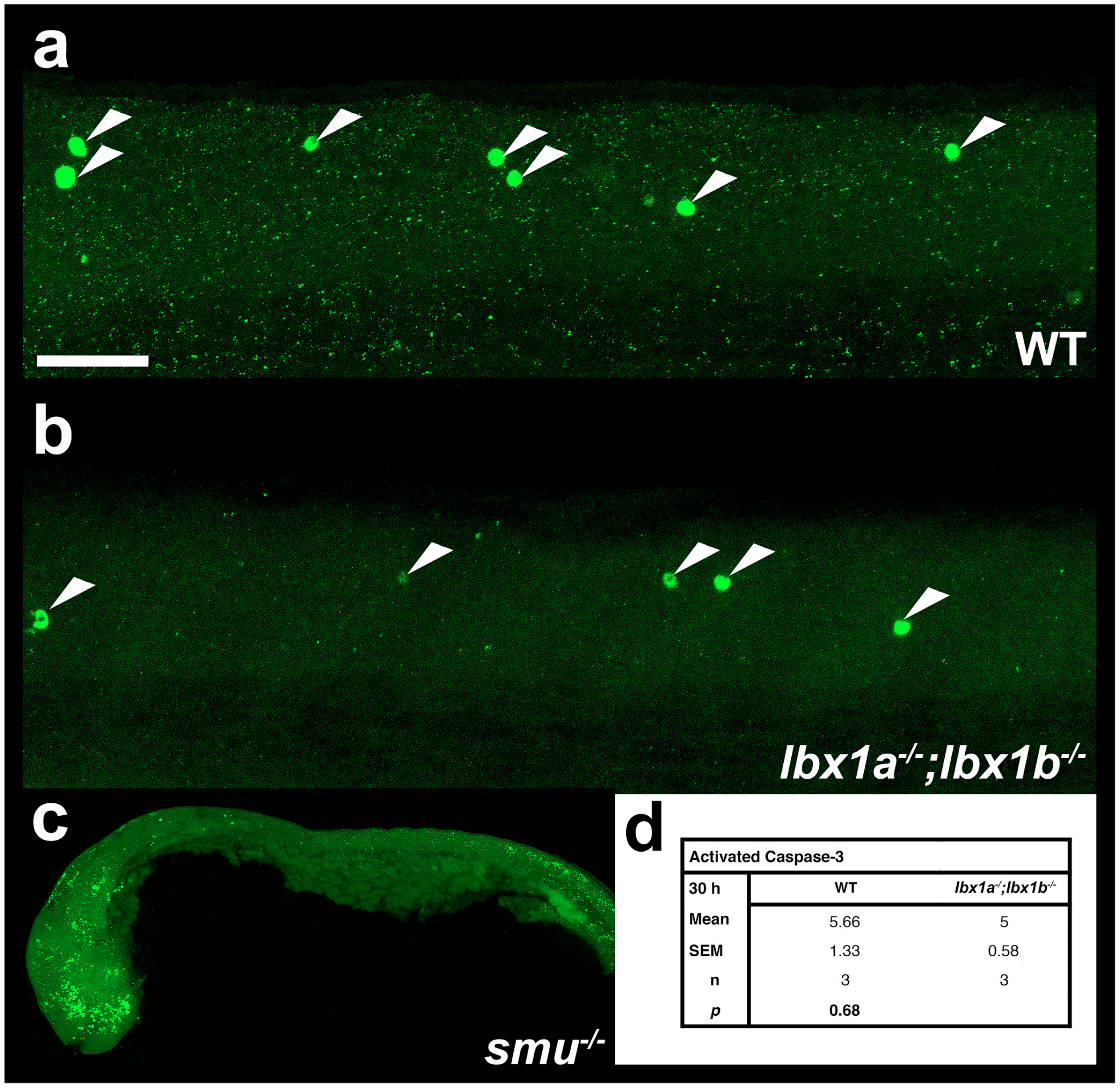
There is no increase in apoptosis in *lbx1a;lbx1b* double mutant spinal cords. **(a-c)** Lateral views of activated caspase-3 immunohistochemistry in zebrafish spinal cord (**a & b**) or whole embryo (**c**) at 30h in **(a)** WT embryo, **(b)** *lbx1a^sa1496^;lbx1b^hu3534^* double mutant and **(c)** *smoothened^b641^* mutant. The latter was used as a positive control as apoptosis is increased in the head and tail regions. In all cases anterior is left, dorsal top, **(a** & **b)** White arrow heads indicate Caspase-3-positive cells. **(d)** Numbers of Caspase-3-positive cells in spinal cord region adjacent to somites 6-10 in *lbx1a^sa1496^;lbx1b^hu3534^* double mutants and WT siblings. Values shown are the mean from 3 different embryos, the S.E.M. and the P value from a student’s t-test. Scale bar = 50 μm **(a & b)** and 200 μm **(c)**.

### Evolution of *lbx* spinal cord expression

To investigate where the differences in *Lbx* spinal expression evolved in the vertebrate lineage, we examined *lbx* gene expression in *Scyliorhinus canicula* and *Xenopus tropicalis*.

Similarity searches in a *S. canicula* (small-spotted catshark) embryonic EST database led to the identification of two *lbx* sequences, unambiguously related to *Lbx1* and *Lbx2* sequences characterized in osteichthyans. We were unable to analyze spinal expression of these genes using *in situ* hybridizations on whole-mount specimens, due to lack of probe penetration into the spinal cord. Therefore, we performed *in situ* hybridizations on embryo cross-sections at stages 25, 28, 31 and 32. Similar to mouse (Cheng et al., 2005; Gross et al., 2002; Kruger et al., 2002; Müller et al., 2002), we observed *lbx1* expression laterally in spinal cord, just above the mid-point of the dorso-ventral axis (Fig. 7a). Interestingly, the putative enhancer region that we used to make our *Tg(1.6Kb lbx1a:EGFP)^SU33^* line is conserved between zebrafish, humans and mouse and is partially conserved in *S. canicula* (Fig. 1), suggesting that this genomic region is at least partly responsible for this conserved spinal cord expression.

**Figure 7.**
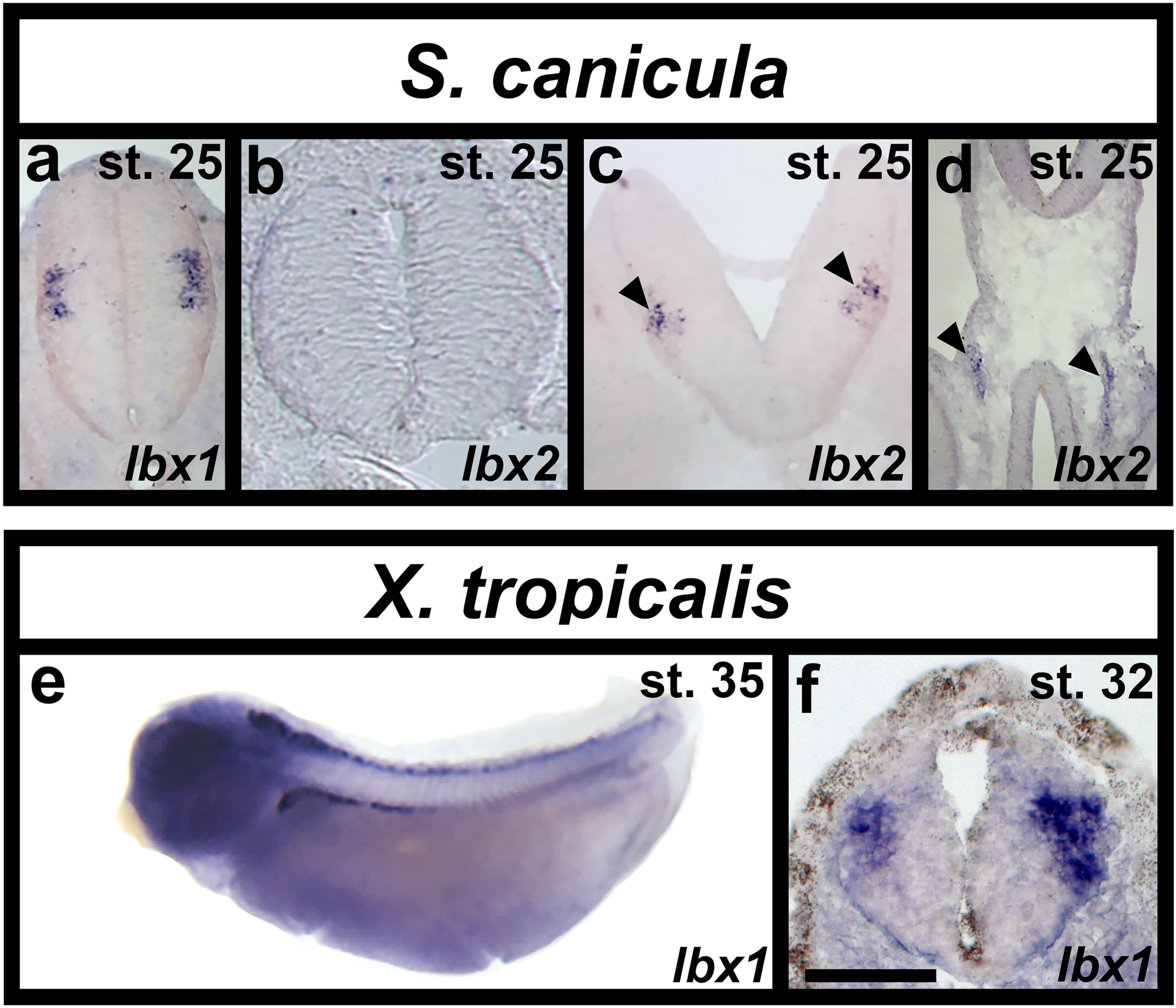
*lbx* expression in *Scyliorhinus canicula* (dogfish) and *Xenopus tropicalis* (frog) spinal cords. **(a-c)** Expression of *Scyliorhinus canicula (S. canicula) lbx1* and *lbx2* in spinal cord and gut (**d**) of cryo-sectioned embryos at stage 25, dorsal top. **(a)** *S. canicula lbx1* is expressed laterally just above the mid-point of the dorsal-ventral axis. **(b)** *lbx2* is not expressed in the spinal cord although it is expressed in the hindbrain (black arrow heads in **c**) and gut (black arrow heads in **d**). **(e)** whole-mount and **(f)** cross-section of *in situ* hybridisation in *Xenopus tropicalis* (*X. tropicalis*) at stages 35 and 32 respectively. **(e)** *lbx1* is expressed in a line of cells along the whole rostral-caudal axis of the spinal cord. **(f)** As in *S. canicula*, *X. tropicalis lbx1* is expressed laterally just above the mid-point of the dorsal-ventral axis of the spinal cord. Scale bar = 140 μm **(a-d)**, 500 μm **(e)** and 50 μm **(f).**

In contrast to *lbx1*, but again similar to mouse (Chen et al., 1999; Moisan et al.), we did not observe any *lbx2* spinal expression at any of these stages (Fig. 7b and data not shown). However, *lbx2* is clearly expressed in both hindbrain (black arrowheads in Fig. 7c) and gut (black arrowheads in Fig. 7d), indicating that our *in situ* hybridization worked.

*Xenopus tropicalis* only has one *lbx* gene, *lbx1* (Wotton et al., 2008). We analyzed expression of this gene from stage 22 to stage 37 (Fig 7e & f and data not shown). *lbx1* is expressed in rostral spinal cord at stage 22 and expression extends more caudally as development proceeds. By stage 35, *lbx1* is expressed along the whole rostral-caudal extent of the spinal cord (Fig. 7e). Similar to small-spotted catshark and mouse, spinal cross-sections show that *lbx1*-expressing cells in *X. tropicalis* are lateral, consistent with them being post-mitotic, and located just above the mid-point of the dorso-ventral axis (Fig. 7f).

## Discussion

*Lbx* genes have crucial functions in mesoderm and nervous system development in a wide range of animals (e.g. Brohmann et al., 2000; Cheng et al., 2005; Gross et al., 2002; Gross et al., 2000; Jagla et al., 1998; Lou et al., 2012; Müller et al., 2002; Schubert et al., 2001). As previously discussed, amniotes have two *Lbx* genes, although only *Lbx1* is expressed in spinal cord (Chen et al., 2001; Chen et al., 1999; Gross et al., 2002; Jagla et al., 1995; Kanamoto et al., 2006; Müller et al., 2002; Schubert et al., 2001). In contrast, zebrafish have three *lbx* genes, as both teleost duplicates of *lbx1* have been retained (Wotton et al., 2008). All three zebrafish *lbx* genes are expressed in spinal cord (Lukowski et al., 2011; Neyt et al., 2000; Ochi & Westerfield, 2009; this report) but before this paper their spinal expression had not been analyzed in detail. Our data show that all three of these genes have distinct spinal expression patterns. Our double-labeling experiments between *Tg(1.6 lbx1a:EGFP)^SU33^* and *dbx2*, and *lbx1a* and *Tg(evx1:EGFP)^SU1^* suggest that zebrafish *lbx1a*-expressing cells are located in the dI6-dI4 spinal region, as the *Tg(1.6 lbx1a:EGFP)^SU33^*-expressing cells are either within the *dbx2* expression domain or slightly dorsal to it, and most of the *lbx1a*-expressing cells are dorsal to *Tg(evx1:EGFP)^SU1^*-expressing cells (Fig. 4; domains often overlap slightly in the smaller zebrafish spinal cord and are not as clearly separated as in amniotes). Finally, our data also demonstrate that a subset of *lbx1a*-expressing spinal cells co-express *pax2a* (which is expressed by V1, V0D, dI4 and dI6 spinal interneurons (Batista & Lewis, 2008)), and many *lbx1a*-expressing spinal cells are inhibitory, whilst a smaller number are excitatory (Fig. 4). Taken together, these analyses suggest that zebrafish *lbx1a* is expressed in dI4, dI5 and dI6 spinal interneurons, like Lbx1 in amniotes. Consistent with this, we previously showed co-expression of *lmx1bb* and *lbx1a*, suggesting that some *lbx1a*-expressing cells are dI5 interneurons (Hilinski et al., 2016).

In contrast to *lbx1a*, *lbx1b* is expressed by progenitor cells, probably in the dP4 or both the dP4 and dP5 domains, as *lbx1b* expression is dorsal to *dbx2* (Fig. 4h) and also dorsal, and medially adjacent, to *lbx1a* (Fig. 3h & i). Consistent with *lbx1b* being expressed in progenitor cells, spinal expression of this gene is almost completely lost in *mindbomb1^ta52b^* mutants (Fig. 2k and l), in which progenitor cells precociously differentiate into post-mitotic neurons (Fig. 2k). This result also suggests that *lbx1b* expression is turned off as cells become post-mitotic, as (in contrast to *lbx2*, see discussion below) there is not even any expanded expression in the caudal spinal cord, where the “youngest” post-mitotic neurons are located at this stage (Fig. 2l; the spinal cord develops in a rostral – caudal gradient). Consistent with *lbx1b* having a different spinal cord expression pattern to *lbx1a*, the CNE that was used to create the *Tg(1.6 lbx1a:EGFP)^SU33^* transgenic line, that recapitulates endogenous *lbx1a* spinal expression, is not found near zebrafish *lbx1b* (Fig. 1). In contrast, the 900bp CNE that we used to create the *Tg(0.9 lbx1a:EGFP)^SU32^* line is conserved between zebrafish *lbx1a* and *lbx1b*. This CNE drives expression in only a very small number of spinal cord cells (Fig. 3g), but there is considerable expression in the hindbrain, where *lbx1a* and *lbx1b* expression is very similar (data not shown).

*lbx2* is expressed in two distinct spinal domains. The ventral domain appears to correspond to progenitor cells located below *Tg(evx1:EGFP)^SU1^*-expressing V0v interneurons (Fig. 4i), suggesting that it is probably the p1 domain, and the dorsal *lbx2*-expressing cells are located in the same dorso-ventral spinal domain as *lbx1a* expressing-cells (Fig. 3 gcsuggesting that *lbx2* may be expressed briefly in some dI4, dI5 or dI6 interneurons or the progenitor cells that give rise to them, although we did not observe any co-expression of *Tg(1.6 lbx1a:EGFP)^SU33^* and *lbx2*. Analyses of *lbx2* expression in spinal cross-sections suggest that some of the more dorsal *lbx2*-expressing cells are post-mitotic, whereas others are located between the progenitor and post-mitotic domains (Fig. 2o and data not shown). Expression of this gene in *mindbomb1^ta52b^* mutants, suggests that *lbx2* is predominantly expressed in progenitor cells, as most of its spinal expression is lost in *mindbomb1^ta52b^* mutants (Fig. 2q). However, there is some expanded expression of *lbx2* in the caudal spinal cord (Fig. 2r) where more recently differentiated spinal cells are located, suggesting that *lbx2* expression persists into some post-mitotic cells, but is turned off relatively quickly after the cells become post-mitotic.

To understand how *lbx* spinal expression has evolved and, in particular, to investigate whether spinal expression of *lbx2* and/or the spinal progenitor domain expression of *lbx1b*, have been gained in the ray-finned lineage or lost in the lobe-finned lineage, we examined expression of *lbx1* and *lbx2* in the small-spotted catshark *Scyliorhinus canicula* and *lbx1* in the African clawed frog *Xenopus tropicalis* (*X. tropicalis* does not have a *lbx2* gene). *S. canicula* is ideally placed to distinguish between ancestral and derived characteristics, as it is a member of the *chondrichthyes* (cartilaginous fishes), which as the sister group to *osteichthyes* (bony fish) provides an outgroup to major *osteichthyan* taxa (Coolen et al., 2008). Our results show that *lbx1* expression in *S. canicula* and *X. tropicalis* is similar to zebrafish *lbx1a* expression (cf Fig. 2 & Fig. 7) and to mouse Lbx1 (Gross et al., 2002; Jagla et al., 1995; Müller et al., 2002; Schubert et al., 2001). In all these species *Lbx1* is expressed in lateral cells just above the dorsal-ventral mid-point of the spinal cord. All together, these data suggest that *Lbx1/ lbx1a* spinal expression is conserved in all vertebrates. However, in contrast, as Lbx1 is not expressed by spinal progenitor cells in amniotes, *S. canicula* or *X. tropicalis, lbx1b* spinal expression was presumably acquired in the teleost lineage after the teleost duplication of *lbx1* into *lbx1a* and *lbx1b*. Our data also suggest that *lbx2* spinal expression was acquired in the ray-finned lineage, as this gene is not expressed in the spinal cord of either amniotes or *S. canicula*. Consistent with the distinct expression patterns of Lbx2 in different vertebrates, our previous analyses did not detect any CNEs in the vicinity of Lbx2 (Wotton et al., 2008; Wotton et al., 2010). In future studies it would be interesting to examine expression of *lbx2* in other teleosts and other extant vertebrates in the ray-finned lineage such as paddlefish, to determine more precisely when the *lbx2* spinal expression domain evolved. One intriguing possibility is that the spinal cord expression of *lbx2* in zebrafish reflects a caudal extension of the hindbrain expression that is seen in *S. canicula*, although interestingly, *Lbx2* is not expressed in the amniote hindbrain (Chen et al., 2001; Chen et al., 1999; Kanamoto et al., 2006).

The fact that both *lbx1a* and *lbx1b* have been maintained in zebrafish and other teleosts suggests that either Lbx1 functions have been subdivided between these two genes or that one or both of them have acquired novel function(s). The observation that *lbx1b* is expressed in different cells to *lbx1a* might suggest the latter. However, our mutant studies suggest that both *lbx1* genes are required for the correct number of inhibitory spinal interneurons, although interestingly only *lbx1a* is required for the spinal cord to have the correct number of excitatory spinal interneurons (Fig. 5). These data suggest that *lbx1b* and *lbx1a* are both required, presumably in succession (given that *lbx1b* is expressed by progenitor cells and *lbx1a* is expressed by post-mitotic cells), for correct specification of dI4 and dI6 interneurons. It also suggests that the specification of inhibitory fates and the inhibition of excitatory fates are regulated by independent mechanisms, with different requirements for Lbx1b function. One possible explanation for this, would be if the acquisition of excitatory fates occurs after the loss of inhibitory fates, and the influence of Lbx1b does not persist long enough to affect the former. While some of the analyses in mouse have focused on Lbx1’s role in specifying neurotransmitter phenotypes (e.g. Cheng et al., 2005), others suggest that in the absence of Lbx1, dI4-dI6 cells transfate into dI1-dI3 interneurons (Gross et al., 2002; Müller et al., 2002). This would also cause a reduction in inhibitory interneurons and an increase in excitatory interneurons as dI4 and dI6 interneurons are inhibitory whereas dI1, dI2, dI3 and dI5 interneurons are excitatory. In this case, the change in cell fate might be a multistep process, with both *lbx1a* and *lbx1b* being required for the early steps, and only *lbx1a* for the latter steps.

The similarity between some aspects of the phenotypes of *lbx1a* and *lbx1b* single and double mutants suggest that post-mitotic *lbx1a*-expressing cells may derive from the *lbx1b*-expressing progenitor domain. Consistent with this, as discussed above, the *lbx1b* expression domain overlaps with the most dorsal *lbx1a*-expressing cells. If *lbx1a*-expressing cells do indeed derive from the *lbx1b*-expressing progenitor domain, this would suggest that these two genes are transiently expressed by the same spinal cells, with *lbx1b* being expressed before *lbx1a*. This would further suggest that some of the cell-type specific regulatory elements that control *lbx1* spinal expression have been retained by *lbx1b* and there has just been a change in the regulation of the temporal specificity of its expression. It would also imply that the more ventral location of many of the *lbx1a*-expressing cells may be due to ventral migration. Interestingly, this would be consistent with mouse, where some of the Lbx1-expressing spinal cells migrate ventrally (Gross et al., 2002).

In conclusion, our data suggest that zebrafish *lbx1a* is expressed by dI4, dI5 and dI6 spinal interneurons and that this expression pattern and the specification of at least these dorsal spinal interneuron populations are conserved in vertebrates. In contrast, *lbx1b* and *lbx2* have novel spinal cord expression patterns that probably evolved in the ray-finned vertebrate lineage (*lbx2*) or in teleosts (*lbx1b*). Our mutant analyses suggest that *lbx1b* and *lbx1a* are required in succession for correct specification of dI4 and dI6 spinal interneurons, although only *lbx1a* is required for suppression of excitatory fates in these cells. Taken together, the data in this paper increase our knowledge of spinal cord evolution and of the genetic mechanisms that establish correct neurotransmitter phenotypes within the spinal cord.

## Acknowledgements

We thank Sophie Lutter for performing preliminary experiments that led to this project. We thank ZFIN for essential zebrafish resources, the Sanger Zebrafish Mutation Project and Stemple lab for providing us with *lbx* mutant alleles, Andrew Waskiewicz for providing us with the *Tg(lbx1b:EGFP)^ua1001^* transgenic line and Patrick Wincker, Corinne Da Silva and Hélène Mayeur for help in identifying and sequencing the *S. canicula lbx* sequences. We are grateful to Adrian McNabb, Tomasz Dyl, Henry Putz, Jessica Bouchard, Paul Campbell, Annika Swanson and Leslie Vogt and several SU undergraduate fish husbandry workers for help maintaining zebrafish lines. We also thank the Marine Biological Association of the United Kingdom, Plymouth and Clare Baker’s lab for providing us with *Scyliorhinus canicula* egg cases and Jim Smith’s Lab for providing us with *Xenopus tropicalis* embryos. This research was funded by NIH NINDS R01:NS077947 and NSF IOS 1755354.

